# Elevated body mass index in youth is associated with dysregulated surrogate markers of neural inhibition & excitation, and internetwork functional dysconnectivity

**DOI:** 10.1101/2024.04.18.590064

**Authors:** AC Reichelt, E Daskalakis, J Cohen, KG Solar, M Saberi, M Ventresca, M Ali, R Zamyadi, SE Scratch, J Hamilton, BT Dunkley

**Affiliations:** Department of Physiology and Pharmacology, Western University, London, ON, Canada; Faculty of Health and Medical Sciences, University of Adelaide, Adelaide, SA, Australia; Institute of Medical Science, University of Toronto, Toronto, ON, Canada; Neurosciences & Mental Health, The Hospital for Sick Children Research Institute, Toronto, ON, Canada; Pharmacology & Toxicology, University of Toronto, Toronto, ON, Canada; Department of Psychology, University of Toronto, Toronto, ON, Canada; Bloorview Research Institute, Toronto, ON, Canada; Department of Paediatrics, University of Toronto, Toronto, ON, Canada; Rehabilitation Sciences Institute, University of Toronto, Toronto, ON, Canada; Department of Endocrinology, Hospital for Sick Children, Toronto, ON, Canada; Department Diagnostic & Interventional Radiology, Hospital for Sick Children, Toronto, ON, Canada; Department of Medical Imaging, University of Toronto, Toronto, ON, Canada; Department of Psychology, University of Nottingham, Nottingham, UK

**Keywords:** Magnetoencephalography, body mass index, childhood and adolescence, paediatric brain imaging, functional connectivity, neural oscillations, lean, overweight, obesity

## Abstract

The developing child and adolescent brain is thought to have an increased vulnerability to the negative impact of obesity and excessive consumption of hyperpalatable and energy-rich foods. In this study, we investigated the neurophysiological effects of overweight and obesity in 30 participants spanning childhood and adolescence (8-19 years), using a naturalistic viewing paradigm with *Inscapes,* in a pseudo-resting-state protocol scan with magnetoencephalography (MEG). Subjects were median split on body mass indices (BMI), categorised into two groups comprising: lower <25 kg/m^2^ (n=15) and higher ≥25 kg/m^2^ (n=15). We assessed spontaneous, regional neural function indexed by oscillatory activity, and functional connectivity within and between intrinsic resting brain networks, including the default mode network, dorsal and ventral attention, somatomotor, visual, language, central executive and salience networks. Elevated BMI was associated with significant reductions in activity of the posterior dominant rhythm, and gamma hyperactivity across widespread cortical areas, suggesting intrinsic neuronal hyperexcitability and disinhibition in children and adolescents. Additionally, we observed low-frequency theta hypoconnectivity between resting state networks including the salience, visual, and default mode networks, and overall reduced global efficiency in brain network structure, suggesting reduced effectiveness in neural communication. These findings underscore the neural impact of body composition on the developing brain, suggesting deleterious alterations in excitation and inhibition from surrogate neural markers associated with neurochemistry and brain networks linked with cognitive and behavioural functioning. These alterations may contribute to the persistent behavioural rigidity and difficulties in adopting healthier eating behaviours into adulthood.

## Introduction

The excessive consumption of ultra-processed, palatable, high fat and high sugar calorie dense foods is the leading cause of obesity. Obesity is not only associated with metabolic abnormalities due to the accumulation of excess adiposity, but also with negative impacts to neural function and cognition (Bocarsly et al., 2015; Greenwood & Winocur, 2005). The prevalence of overweight and obesity in youth has risen dramatically worldwide, from just 4% in 1975 to over 18% in 2016 (Keating et al., 2014). In developed countries, approximately 20% of children and adolescents are currently classified as overweight or obese (Mitchell et al., 2009; Ng et al., 2014).

Obesity has been associated with changes in brain structure (Dekkers et al., 2019), and during childhood there are profound physiological and neurological health effects, with young, immature brain being particularly vulnerable to the negative functional consequence of obesity and overconsumption of hyperpalatable, ultra-processed calorie-dense foods (Lowe et al., 2020; Reichelt & Rank, 2017).

Neuroimaging studies have demonstrated that the frontal cortices of the brain are the final regions to reach maturity (Giedd, 2004), with the continued development of the frontal cortex during adolescence driven by heightened plasticity and increased myelination of neural circuits, which can be modulated by environmental influences, including diet (Baker & Reichelt, 2016; Lowe et al., 2019; Reichelt, 2016).

The young brain has immature self-regulatory systems combined with mature reward-driven motivational systems, which may cause difficulties in limiting consumption of highly palatable processed foods (Lowe et al., 2020; Reichelt & Rank, 2017). In tandem with the negative metabolic impacts, the adverse neurological effects of obesity may be exacerbated by the heightened window of neuroplasticity in youth which leads to enduring changes to motivating behaviours and cognitive function (Lowe et al., 2020). In particular, the corticolimbic pathway connecting the medial prefrontal cortex (mPFC) and the amygdala undergoes dramatic structural reorganisation during adolescence (Casey et al., 2015; Jalbrzikowski et al., 2017; Silvers et al., 2017), which is associated with behavioural rigidity following obesogenic diet consumption (Labouesse et al., 2017; Vega-Torres et al., 2022).

Overweight and obesity, especially during childhood and adolescence, may therefore present an enhanced risk to brain function. In particular, the fine balance between excitation and inhibition in the brain is believed to play a critical role in brain function. Rodent models of diet-induced obesity have shown reductions in inhibitory neurons and inhibitory neurotransmission in the frontal cortices (Baker & Reichelt, 2016; Reichelt et al., 2019; Sandoval-Salazar et al., 2016; Seabrook et al., 2023; Thompson et al., 2017). As such, excitatory and inhibitory balance within the frontal cortex is influenced by the deleterious presence of obesity - however, this idea has not yet been examined in young people of differing BMI.

In this study, we used magnetoencephalography (MEG), a non-invasive neuroimaging technique, to directly study neural function and determine the impact of BMI on neural activity and network function in children and adolescents. A key component of neural activity are neural oscillations, or ‘brain waves’. Neural oscillations are rhythmic fluctuations in neuronal excitability that allow ensembles of neurons to process information, a core mechanism of neuronal circuits that supports cognitive, perceptual and emotional processes. MEG has been used to measure spontaneous neural activity through development and in neuropsychiatric conditions, characterising typical and pathological neural circuits (Hunt et al., 2019; Wiesman et al., 2022).

Regional oscillatory activity can be used as proxies for excitation and inhibition in the brain. For example, alpha band activity (8–14 Hz) is thought to represent localised neural inhibition (Jensen & Mazaheri, 2010) and is driven by strong and reciprocal thalamocortical connections (Roux et al., 2013). Conversely, gamma band activity (∼30-150 Hz) is generated by the interplay between neural excitation and inhibition (Orekhova et al., 2018), and primarily cortical microcircuits involving inhibitory interneurons, such as parvalbumin (PV) cells (Sohal et al., 2009).

The synchronisation of neural oscillations *between* brain regions can also be quantified to measure functional connectivity (Fries, 2015). Functional connectivity is a surrogate measure of the brain’s ability to communicate and coordinate information. In this regard, there are a number of important brain networks that take on particular roles. The default mode network (DMN) is involved in coordinating network dynamics (de Pasquale et al., 2012) and supporting spontaneous cognition, self-reflective thought or attention to internal stimuli (Buckner et al., 2008). Other key networks include the Central Executive Network (CEN) and the salience network.

The CEN is a large-scale brain network involved in a variety of cognitive functions, including working memory, attention, and decision-making (Sherman et al., 2014). It is also involved in the coordination of other brain networks and the integration of information from different sources. The salience network focuses attention, motivates behaviour, and processes emotions (Seeley, 2019), detecting important stimuli and coordinating brain activity across networks, with dysfunction within and between these networks linked to psychiatric disorders (Menon, 2011).

Resting-state functional MRI (fMRI) studies have revealed altered DMN activity in adults with obesity (Doucet et al., 2018; Stice & Burger, 2019; Tregellas et al., 2011), and adults at risk of developing obesity based on familial heritage (Mahmoud et al., 2022). Furthermore, a single pilot study used MEG to examine alterations in functional connectivity in 11 female adolescents with severe obesity compared to controls, and observed increased resting-state functional connectivity, indicating hyperconnected functional coupling between cortical areas in obesity (Olde Dubbelink et al., 2008). However, no studies have examined functional connectivity specific brain networks or the topological structure of these networks in children and adolescents with overweight and obesity.

Here, we investigated the neurophysiological effects of higher vs lower body mass index (BMI) in 30 participants spanning childhood and adolescence (8-19 years), using a naturalistic viewing paradigm with *Inscapes,* in a pseudo-resting-state MEG. Participants were categorised into two groups based on a median split of BMI across the group: lower <25 kg/m^2^ and ≥25 kg/m^2^. We employed the DISCOVER-EEG biomarker discovery pipeline for clinical neuroscience, which recommends a battery of neural markers for examining neurophysiological functioning in clinical research (Gil Ávila et al., 2023). We predicted that those with higher BMI group would exhibit decreased alpha activity, a marker of cortical disinhibition and thalamocortical dysfunction, and gamma hyperactivity, a marker of neural excitability and PV functioning. Moreover, we predicted network-level alterations in functional connectivity and topological graph structure, including reduced global efficiency, clustering and small worldness.

## Methods

### Participants

Thirty youth participants were recruited with ages spanning childhood and adolescence (8-19 years). Height and weight were used to calculate each participant’s body mass index (BMI, body mass in kilograms / height in metres^2^) and participants were stratified using a median split into 2 groups, n=15 with a low BMI<25 (mean age=14 years, std=2.97 years, 8-19 years range, 8 males) and n=15 with a high BMI≥25 (mean age=12.9 years, std=2.65 years, 8-18 years, 8 males).

Inclusion criteria for this study included male and female children and adolescents aged 6-19 years with the ability to understand task instructions in English. Exclusion criteria included those with a diabetes diagnosis, history of drug use or alcoholism, smoking, major head trauma, history of non-removable metal in the body (e.g., medical devices, major orthodonture, etc), those over 137 kg (due to scanner bed weight limits), and those given prescription of glucocorticoid medications. All participants gave informed consent. Research ethics approval was granted by the Hospital for Sick Children Research Ethics Board.

### Procedure and MEG data acquisition

Participants were directed to the magnetically shielded MEG scanner room where a 5-minute eyes-open *Inscapes “*pseudo-resting-state” scan was collected (151 Channel@600 Hz; CTF System, Coquitlam, Canada). Resting state MEG measures spontaneous, intrinsic neurophysiological activity and network functional connectivity patterns. During the scan, participants watched the *Inscapes* video, a naturalistic viewing paradigm (Vanderwal et al., 2015), comprising a computer generated video and audio landscape, and instructed to “remain in a state of resting wakefulness, stay within their thoughts, and allow their mind to wander.” *Inscapes* has been shown to recapitulate resting state network activity analogous to ‘traditional’ fixation cross resting state, and is also known to increase participant compliance / tolerance with scan parameters and subsequently reduce the amount of head motion artefacts (Vandewouw et al., 2021). Continual head position localisation was captured through fiducials attached to the nasion and left and right pre-auricular point in each participant.

### MEG data analysis

MEG data was analysed with the FieldTrip toolbox (Oostenveld et al., 2011). The continuous, resting state data was first filtered using a 4th order Butterworth band-pass filter with a lower limit of 1 Hz and an upper limit of 150 Hz. Additionally, notch filters were applied to remove the main line interference at 60 Hz, along with the harmonic at 120 Hz. The data was divided into 10 second epochs following inspection of the head motion data, with epochs being excluded if the head position deviated more than 10 mm from the starting head position, and head velocity was more than 5 mm/s.

In order to remove cardiac and eye movement/blink artefacts, Independent Component Analysis was performed (using the ‘fastica’ algorithm in FieldTrip), following which, visual inspection of components was performed and components corresponding to cardiac or ocular artefacts were removed manually (by an experienced analyst). The post-ICA cleaned data was used for subsequent analysis.

In order to localise cortical sources of activity we defined 78 cortical regions defined by the Automated Anatomical Labelling Atlas (AAL) (Tzourio-Mazoyer et al., 2002). Additionally, 54 node coordinates were used to define resting state networks, including the DMN, dorsal and ventral attention, central executive, visual, motor, salience and language networks (de Pasquale et al., 2012). A linearly constrained minimum variance (LCMV) beamformer (Van Veen et al., 1997) was employed to estimate electrophysiological activity at these predetermined ROI locations within the brain. MEG data co-registration was performed by creating a single-shell head model to create the forward solution for each individual participant using an age-appropriate anatomical T1-weighted template (Evans & Brain Development Cooperative Group, 2006). A common spatial filter was calculated for each of seed locations, and a covariance matrix was generated (regularised with a 5% Tikhonov regularisation factor) using the epoched, artefact free sensor level data. Finally, source level, broadband time series were calculated for the centroid of each of the ROIs (Sekihara et al., 2001).

Source level time-series data was used to calculate the regional power spectral density (PSD) for each of the 78 cortical regions. The time-series data for each region was z-score normalised (mean centred, variance normalised) and the PSD was calculated for each epoch using the Welch method; which were in turn separated into canonical frequency bands: Delta (1-3 Hz), Theta (4-7 Hz), Alpha (8-14 Hz), Beta (15-30 Hz), Low Gamma1 (30-55 Hz), Low Gamma2 (65-80 Hz), and High Gamma (80-150 Hz). The PSD values for each frequency band were averaged across epochs for subsequent analysis.

For the functional connectivity analysis, broadband, source level time-series data for the 54 cortical regions were filtered into the aforementioned canonical frequency bands. Amplitude Envelope Coupling (AEC) was chosen as the connectivity measure of choice due to its stability and reliability (Colclough et al., 2015). One issue with AEC connectivity is the leakage between source signals at separate locations (Brookes et al., 2012), with leakage correction applied with the multivariate symmetric orthogonalization procedure (Colclough et al., 2015). Following leakage correction, the analytical signals were computed for each frequency band using the Hilbert transform resulting in the amplitude envelope of the source time series data. Pearson correlation coefficients were calculated for each possible pair-wise combination of the 54 regions resulting in an 54 x 54 adjacency matrix.

### Statistical approach

Inferential tests for between-groups differences in outcome measures (e.g. age, heigh, weight) were conducted in JASP using either an independent samples Student’s t-test for normally distributed data with equal variance, or Mann-Whitney U test when distributions were non-Gaussian.

To compare group differences in regional PSD, non-parametric Wilcoxon rank-sum tests were performed for each of the 78 cortical regions, for each frequency band. The resulting p-values were corrected for multiple comparisons across each region using the Benjamini-Hochberg correction for false discovery rate (FDR-BH; Benjamini et al., 1995), with p < 0.05.

To identify functional connectivity patterns that were significantly different between groups, adjacency matrices were concatenated for each group, and averaged across each network. Given the unknown nature of distributions, non-parametric permutation testing was performed to estimate the p-value (10,000 iterations), with a false discovery rate correction for multiple comparisons, p < 0.05. Graph theory measures of network topology were computed with the Brain Connectivity Toolbox (BCT) (Rubinov & Sporns, 2010). Finally, BrainNet Viewer was used for visualisation purposes of regional oscillatory activity and connectivity (Xia et al., 2013).

## Results

### Participant characteristics

There was no significant difference in mean age between the BMI≥25 (mean age = 12.9 years, std = 2.65) and the BMI<25 group (mean age = 14 years, std = 2.97, t(28) = -1.1, p = .28), and no significant difference in mean height between the BMI≥25 (mean height = 1.65m, std = 0.17) and the BMI<25 group (mean height = 1.59m, std = 0.15; U(28) = 74, p = .12). There was a significant difference in overall mean mass between the BMI≥25 (mean mass = 88.41 kg, std = 20.1) and the BMI<25 group (mean mass = 44.19 kg, std = 12.06; t(28) = -6.8, p < .001).

### Elevated BMI is associated with reduced posterior dominant rhythm power

Examining averaged lobewise power spectrum curves from the virtual sensors placed at AAL node locations, both groups exhibit characteristic aperiodic 1/f power profiles (**Figure 1A**), with pronounced periodic activity in the alpha range (∼8-12 Hz) and beta range (∼15-25 Hz), especially in parieto-occipital and fronto-temporal regions, respectively. Those in the BMI≥25 group show marked reductions in apparent peak alpha power in posterior regions (∼10 Hz), as well as apparent increases in broadband gamma activity (+30 Hz) across multiple lobes.

**Figure 1.**
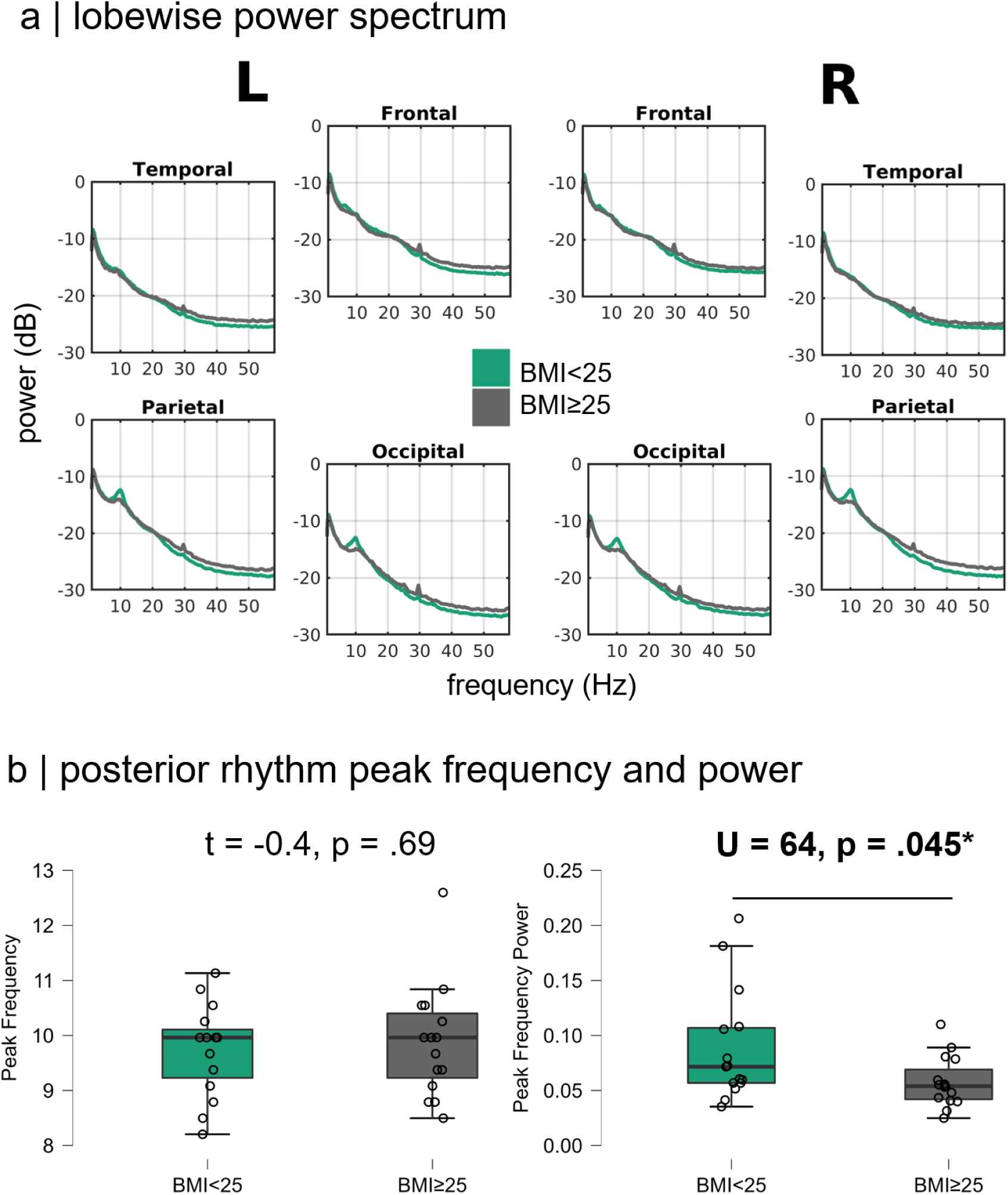
Spectral profiles and posterior dominant rhythms by BMI. **A.** Lobewise spectral power plots show both groups present with typical 1/f aperiodic power spectrum curves, and periodic activity at alpha and beta frequencies, with cortical-wide increases in broadband gamma activity, and reductions in peak alpha amplitude, the latter evident in parieto-occipital areas. **B.** Peak posterior rhythm did not differ between groups, but peak frequency maximum power was significantly lower in the BMI≥25 group.

We tested the peak frequency of the dominant posterior rhythm and power of the individual peak frequency between groups, extracted from the ‘occipital’ grouping of seeds from the AAL atlas. There was no significant difference in mean peak frequency between the BMI≥25 group (mean peak freq = 9.89 Hz, std = 1.03 Hz) and the BMI<25 group (mean peak freq = 9.75 Hz, std = 0.83 Hz; t(28) = -0.4, p < .69; **Figure 1B**). There was a significant group difference in peak frequency maximum power between the BMI≥25 group (mean peak power = 0.058 dB, std = 0.023 dB) and the BMI<25 group (mean peak power = 0.089 dB, std = 0.051 dB; U = 64, p = .045; **Figure 1B**).

### Elevated BMI is associated with cortical wide broadband gamma hyperactivity

The BMI≥25 group exhibited significantly increased spontaneous broadband gamma activity, across numerous band-limited ranges, including 30-55 Hz, 65-80 Hz, and high gamma 80-150 Hz ranges (p < .05, FDR-corrected). These differences occurred across every lobe of the brain at low gamma (30-45 Hz) ranges (**Figure 2A**), with a similar topography for gamma in the 55-80 Hz range, and a smaller number of regions showing significant differences at the high gamma (80-150 Hz) range (**Figure 2B**).

**Figure 2.**
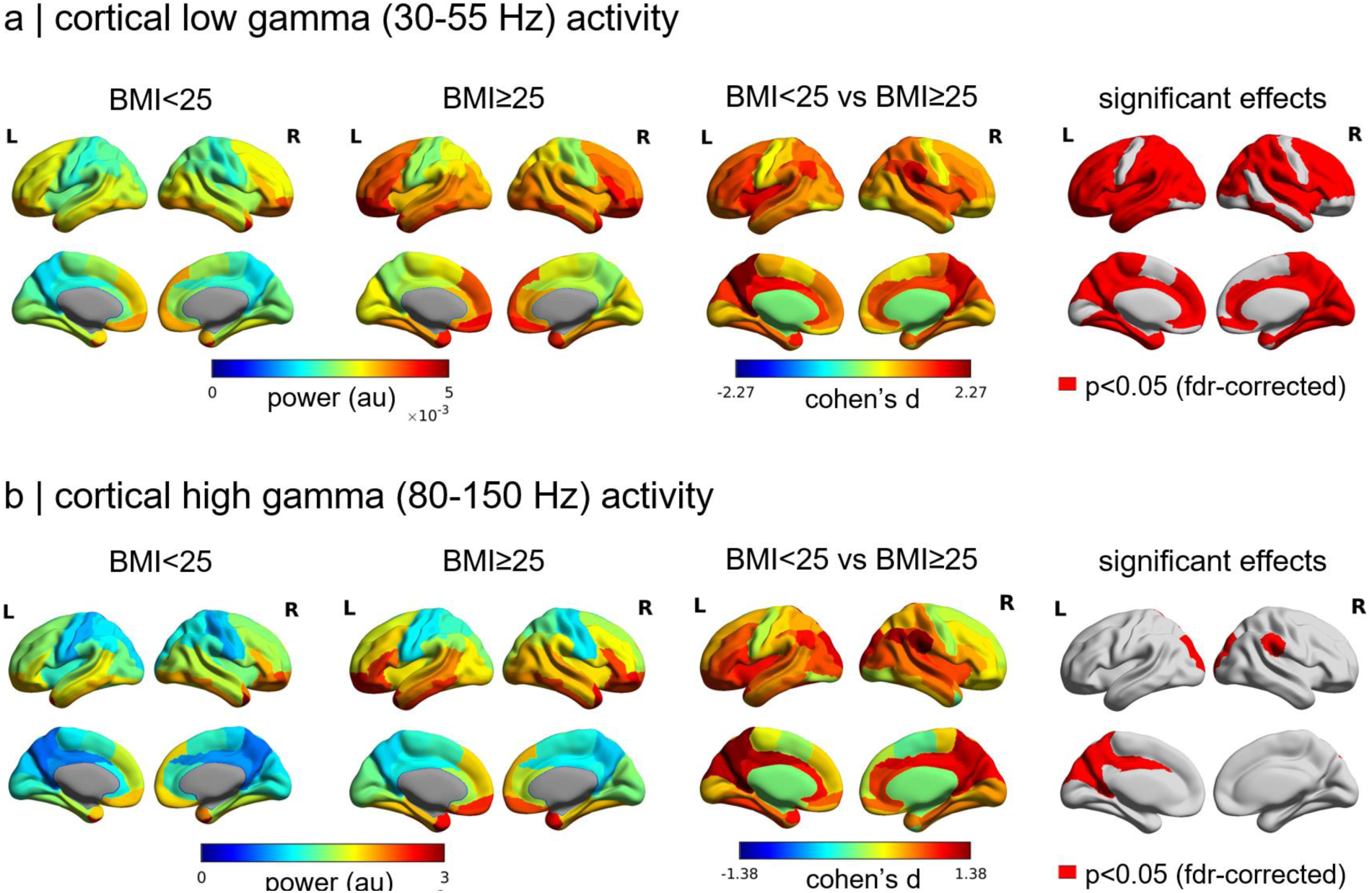
Higher BMI is associated with cortical wide broadband gamma hyperactivity. **A.** Cortical oscillatory activity in the low gamma band (30-45 Hz) was significantly increased in the BMI≥25 group across widespread areas of the cortex, with the greatest effect noted in medial parietal areas. **B.** In the high gamma band (80-150 Hz) similar effects were noted in the medial parietal areas, as well as the right temporo-parietal junction.

Regions with the greatest effect sizes defined by Cohen’s d include the posteromedial cortex, including the medial parietal and precuneus areas, as well as dorsal visual stream areas, with a consistent significant effect in the right temporo-parietal junction for all three gamma bands (with effect sizes of d = 2.27 for gamma 30-45 Hz, through to d = 1.38 for high gamma 80-150 Hz). No significant effects were noted for any of the other frequency bands analysed.

### Those with higher BMI exhibit functional dysconnectivity between numerous resting state networks and decreased global efficiency

Significant differences in theta-mediated functional connectivity were also observed in between group comparisons (p < 0.05, FDR-corrected), with the BMI≥25 group showing reduced connectivity within the visual network, and for intranetwork connectivity for the visual-to-dorsal and ventral attention networks, and the DMN (**Figure 3A**). Dysconnectivity was also noted for the DMN-to-salience and language networks, and the Salience-to-central executive and ventral attention networks. No significant differences were observed for any other frequency band analysed (p > 0.05, FDR-corrected).

**Figure 3.**
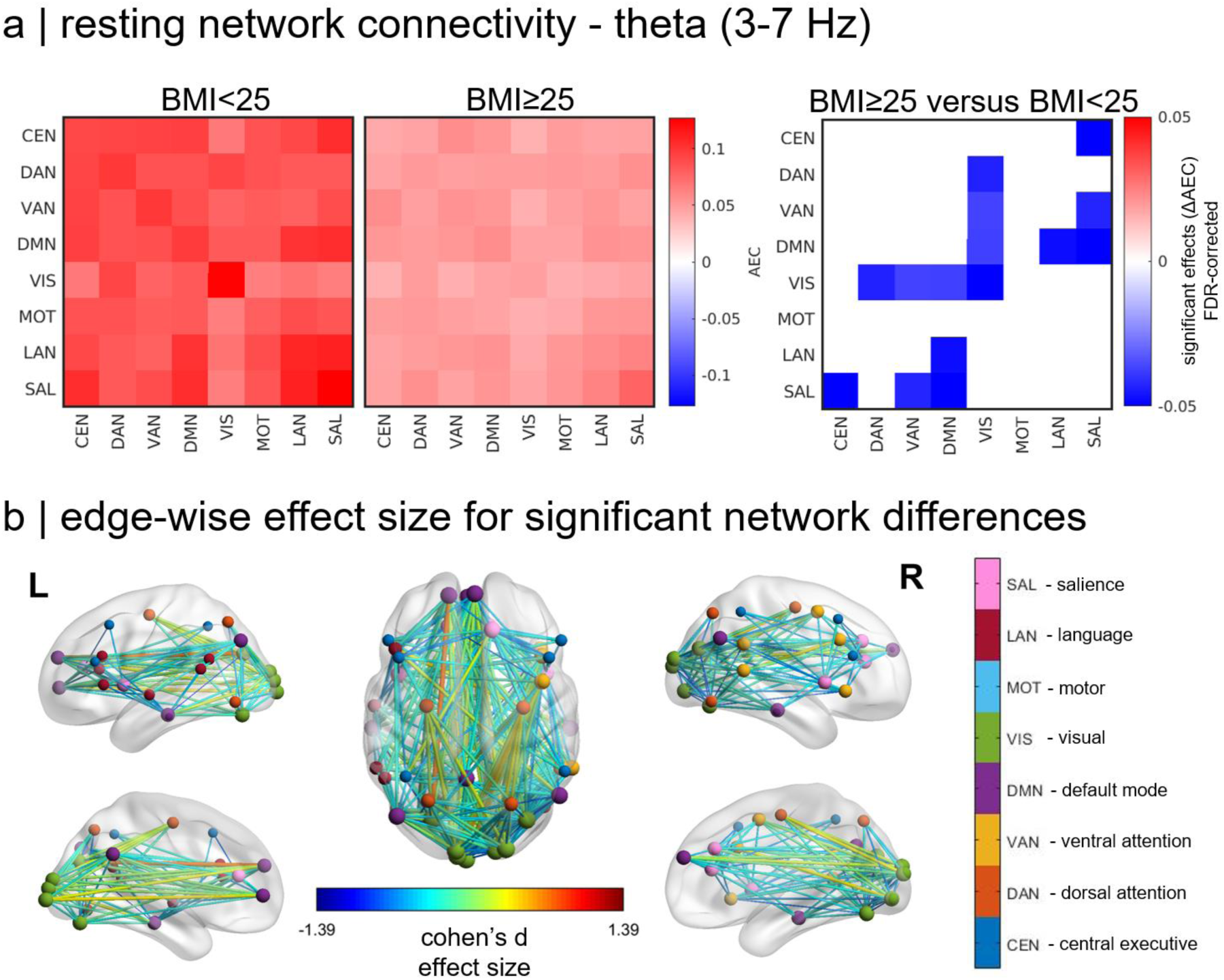
Resting state network dysconnectivity mediated by theta oscillations is present in higher BMI. **A.** Theta functional connectivity was reduced in the BMI≥25 group compared to the BMI<25 group, especially for default mode network internetwork connectivity to the visual, motor, language and salience networks, and within the visual network, and for visual to dorsal and ventral attention networks. **B.** Edgewise effect sizes displayed in brain space for significant, between group inter and intranetwork tests identified in **A** reveal widespread resting network reductions in connections across many areas of the brain.

Plotting edgewise effect sizes (Cohen’s d) of the significant network-level effects above show the majority of connection weights were reduced in the BMI≥25 group, with only a small percentage of connections showing elevated connectivity compared to the BMI<25 group (**Figure 3B**).

Subsequently, the entire functional connectivity matrix (e.g. all intra and interwork network edges) for the theta mediated AEC were tested for between group differences in overall network topology, including global efficiency (GE), clustering coefficient (CC), and small worldness (SW; **Figure 4**). There was a significant difference in global efficiency between the BMI≥25 group (mean GE = 0.098, std = 0.021) and the BMI<25 group (mean GE = 0.127, std = 0.044; U = 52, p = .011; **Figure 4A**). There were no significant differences between groups in either clustering coefficient (**Figure 4B**) or small worldness (**Figure 4C**).

**Figure 4.**
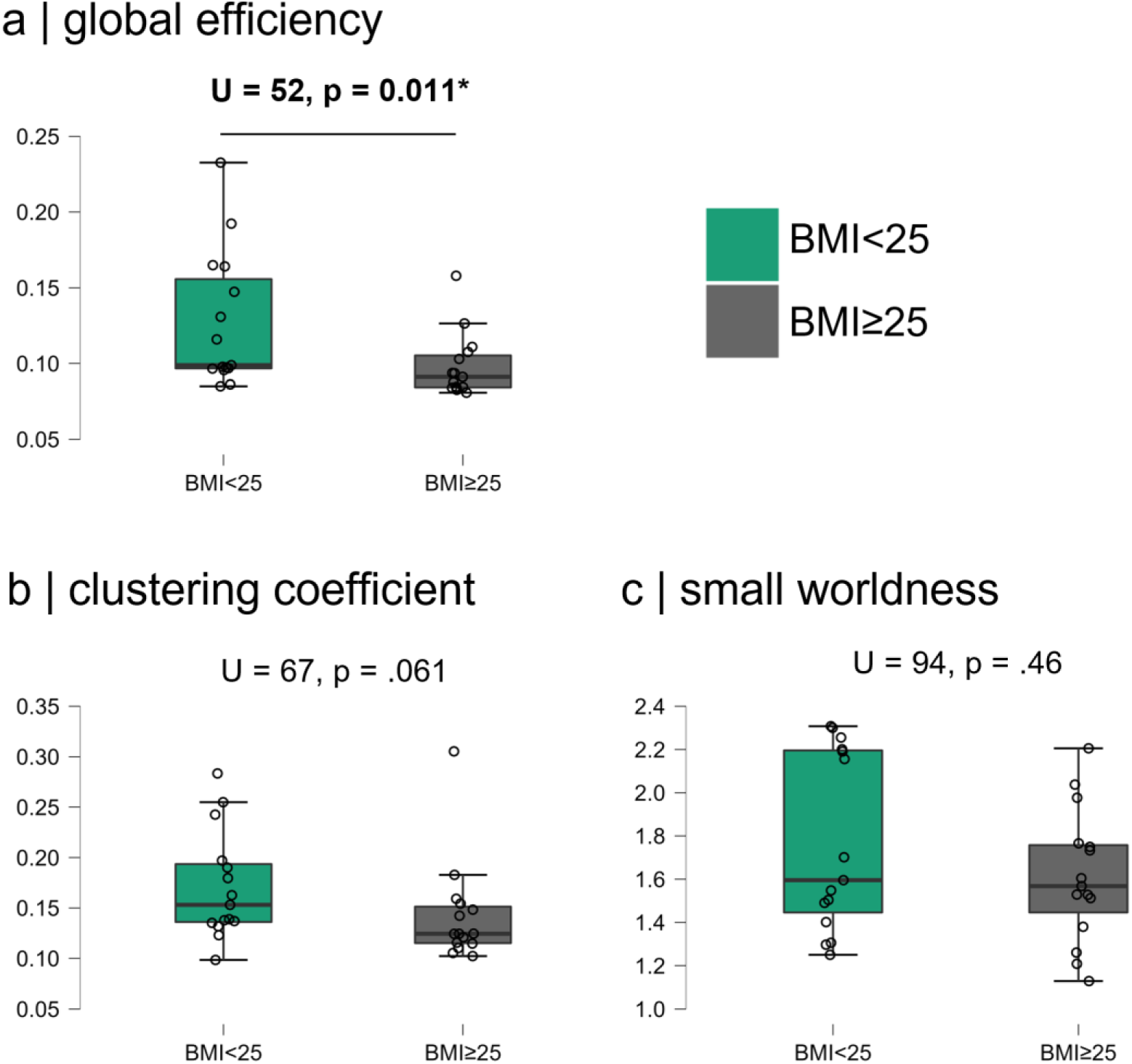
Topological graph measures for theta functional connectivity. Global network graph structure of theta-mediated functional connectivity shows significant reductions in **A.** global efficiency in the BMI≥25 group, with no significant differences in **B.** clustering coefficient or **C.** small worldness.

**Figure 5.**
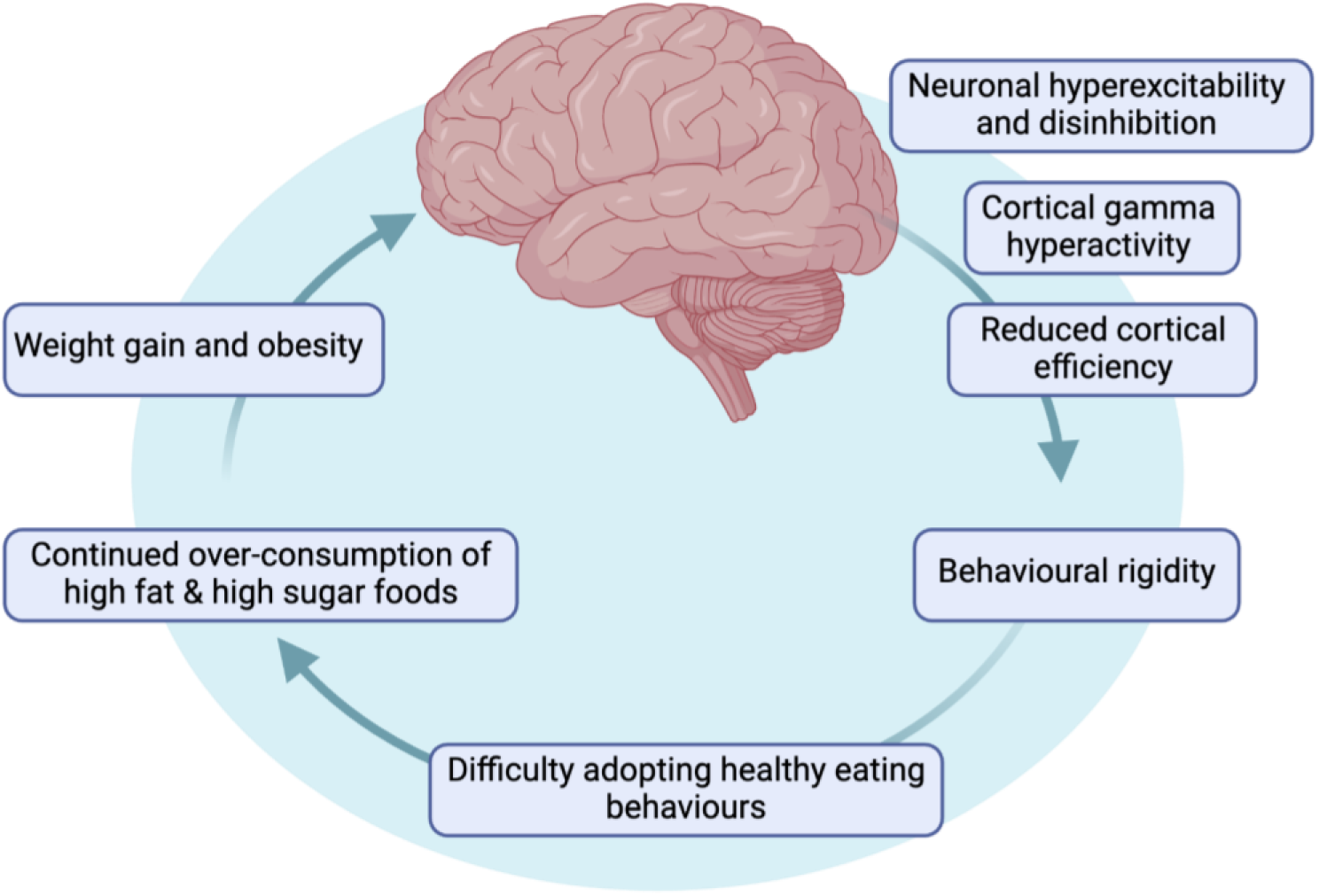
Multiscale mechanisms underlying the closed loop potentiation of obesogenic behaviours and impact on the brain. Figure drawn in Biorender.com.

## Discussion

### Summary

We report the first evidence of dysregulated neurophysiological functioning, consistent with altered neural excitation and inhibition, and changes in resting state brain networks measured with MEG in children and adolescents with elevated BMI. We observed significant alterations of neural functioning in frequencies associated with intrinsic neural inhibition (alpha activity) and excitation (gamma activity). These effects were most prominent in the posteromedial cortices, including the visual / occipital and medial parietal regions, specifically, the precuneus and posterior cingulate cortex, key brain hub areas involved in the fronto-parietal / CEN, a network that serves cognitive control and mental flexibility mechanisms. We also observed reduced functional connectivity between multiple brain networks, including the visual network, visual-to-dorsal and ventral attention networks, and the DMN. Dysconnectivity was also noted for the DMN-to-salience and language networks, and the Salience-to-central executive and ventral attention networks.

Dysregulation in these areas may underlie some of the cognitive deficits that are observed in those with higher weight, cognitive issues that could precipitate and/or exacerbate the development of severe psychiatric conditions (Wang et al., 2023). These novel findings demonstrate the neurophysiological impact of obesity during a key window of neurodevelopment.

### Regional high-frequency hyperactivity and low-frequency hypoactivity

We observed neural signatures consistent with disinhibited and hyper-excitable cortical regions in children and adolescents with elevated BMI. Group contrasts revealed significantly elevated gamma, particularly in medial parietal cortices, and the posterior cingulate cortex – key regions for top-down cognitive control and mental flexibility. Synchronisation of neuronal assemblies at gamma frequencies is closely associated with cortical processing and during typical brain functioning, inhibition of excitatory pyramidal cells through GABAergic interneurons lead to the emergence of neural oscillations in the gamma-frequency band (Womelsdorf et al., 2006). Our observation of elevated gamma indicates hyper-excitability in regions critical for cognitive processes, which may be driven by dysregulated inhibitory interneuron populations (Guan et al., 2022).

Elevated gamma might also be implicated in regional inhibitory gating - these results suggest tonic disinhibition in elevated BMI. Previous studies in animal models have shown that injury to GABAergic interneurons, specifically PV-containing interneurons, causes abnormal increases in spontaneous gamma-band activity (Carlén et al., 2012; Del Pino et al., 2013; Korotkova et al., 2010) and is associated with imbalances in the excitatory/inhibitory ratio within neural networks (Buzsáki & Wang, 2012). Alterations in inhibitory GABAergic function have been shown in rodent models of diet induced obesity. Rats fed eight weeks of a high fat diet had decreased GABA levels in the frontal cortex (Sandoval-Salazar et al., 2016), and other studies have shown four weeks of binge-like high fat diet consumption led to a reduction in PV+ interneurons (Baker & Reichelt, 2016; Reichelt et al., 2019), and persistent damage to PV+ neurons in mice with diet-induced obesity following experimental stroke (Pintana et al., 2019).

Disinhibition of excitatory neurons may be caused by dysregulation of PV-containing interneurons either directly or through changes to the perineuronal net (PNN) surrounding these neurons. PNNs are a form of specialised extracellular matrix that envelops subtypes of neurons, most frequently PV interneurons, providing a protective microenvironment and controlling plasticity (Wingert & Sorg, 2021). PNNs tightly regulate PV interneuron excitability (Hanssen et al., 2023), and removal of PNNs decreases the mean spiking activity and increases the spiking variability of PV interneurons (Lensjø et al., 2017). Previous studies have shown reduced PNN staining surrounding PV expressing interneurons consumption of hypercaloric diets (Dingess et al., 2018). PNNs have a delayed developmental trajectory in the frontal cortices (Mauney et al., 2013), potentially rendering PV interneurons in the young brain at risk of damage from the neuroinflammatory milieu generated by obesity (Guillemot-Legris & Muccioli, 2017). Abnormal gamma activities may provide a proxy measurement of GABA-ergic interneuron dysfunction (Huang et al., 2019). Therefore diet-induced disruption of PNNs and PV neurons leading to disinhibition of pyramidal neurons may be a mechanistic process underpinning our observed increases in cortical gamma oscillations.

As predicted, we observed decreased alpha in posterior regions in the BMI≥25 group, further suggesting decreased inhibition in these areas. This observation is in keeping with previous studies using EEG that have described alpha band deviations in children (Schmidt et al., 2018) and adults (Babiloni et al., 2011; Del Percio et al., 2013) with overweight or obesity in eyes-closed, resting-state conditions, but this effect was not shown in an eyes-open condition in people with obesity (Kösling et al., 2022). These alterations in alpha frequency oscillations at resting state may reflect abnormal arousal and vigilance processes in young people with increased BMI, and also supports a potential further effect of high BMI on the dysregulation of inhibitory signalling – potentially leading to increased cortical gamma oscillatory activity.

### Altered network connectivity

Functional connectivity describes the co-varying activity profiles of topographically dispersed brain regions (Biswal et al., 1997) and provides a dynamic picture of brain activity. The patterns of spontaneous neuronal activity are believed to reflect direct and indirect anatomic connectivity (Vincent et al., 2007). We observed numerous networks showing altered functional connectivity in elevated BMI. These networks support executive functions by exerting potent top-down effects that are responsible for manipulating information in working memory to consider possible outcomes to influence critical mechanisms including decision-making, problem-solving, and emotion-regulation. Resting-state functional connectivity offers one way to assess the integrity of communication between functionally coupled brain regions.

Disruptions in connectivity is a common neural signature observed in neurological and psychiatric disorders. Network-based analysis identified significantly decreased connectivity mediated by theta frequency amplitude coupling across a distributed set of regions, including fronto-parietal and temporal areas, and key nodes in the DMN. The DMN showed multiple inter-network alterations, with the DMN containing nodes in the parietal areas and precuneus – critical regions that have key roles in attention and higher-order cognition. This weakened functional connectivity pattern in the DMN with the precuneus and posterior cingulate cortex as the key affected nodes has been associated with cognitive impairment, and it is particularly notable that the precuneus and posterior cingulate cortex play a pivotal role in regulating the internal activities of the DMN (Fransson & Marrelec, 2008). Moreover, decreased network connectivity in the theta frequency may underpin alterations to motivated processes, as resting state theta is related to reward dependent learning (Massar et al., 2014), and is also generated within the hippocampus, a brain region known to be negatively impacted both functional (Barbosa et al., 2023) and structurally by obesity (Cherbuin et al., 2015).

Our findings indicate that increased BMI in young people is associated with alterations in cortical communication processes, demonstrated by decreased global organisation efficiency. A previous study using fMRI demonstrated that people with obesity exhibited significantly reduced global and local network efficiency as well as decreased modularity compared with healthy controls, showing disruption in small-world and modular network structures (Baek et al., 2017). Global brain connectivity identifies “hubs,” which are densely connected and metabolically costly, and have a broad influence on brain function. Obesity in adults has been shown to result in lower global brain connectivity, and favours networks important for external orientation over those monitoring homeostatic state and guiding feeding decisions, which is in keeping with the central and peripheral metabolic dysfunction associated with overweight and obesity (Geha et al., 2017). Therefore, our observed reductions in brain connectivity in youth with elevated BMI may dysregulate cognition and also derail the ongoing development of the brain through adolescence and early adulthood, resulting in predispositions to alterations in cognitive processes and mental health disorders like depression (Clasen et al., 2014). In particular, alterations to activity within the vmPFC are associated with deficits in emotion regulation in individuals with obesity (Steward et al., 2019), and impulsive eating behaviours have been associated with suboptimal performance of the Iowa Gambling Task, a decision making task reliant on vmPFC activity (Steward et al., 2019).

Another possible explanation for our results is the potential difference in brain composition between individuals with obesity and normal weight. Using MRI, Haltia et al. showed increased white matter in obese compared to control subjects in brain regions, including the temporal lobes, which may reflect altered functional communication between regional neuronal populations (Haltia et al., 2007).

However, as we measured BMI as opposed to adiposity, we can only speculate that adiposity varied between groups. Nevertheless, functional connectivity differences observed between our high and low BMI groups may indicate altered white matter tracts, and future studies may utilise diffusion weighted imaging in combination with MEG to further define alterations.

### Limitations and Future Directions

While this study offers valuable insight into the functional connectivity across different frequency bands in children and adolescents with BMI in the overweight and obese range, there are several limitations. Using BMI as a measure of body composition has limitations, particularly in young people. The interpretation of BMI in children and adolescents in terms of the classification of obesity and overweight depends on weight, height, age, and sex, whereas for adults, BMI depends only on height and weight. In relation to our current study population, we matched subject groups by age, sex and height as much as possible. Moreover, a BMI greater than the 85th percentile in children and adolescents is considered overweight, and generally reflects a BMI ≥ 25kg/m^2^ within our study sample age range based on CDC classification (Kuczmarski et al., 2002).

In general, BMI measurements cannot differentiate between fat and fat-free mass. Therefore, an elevated BMI may not reliably reflect the accumulation of adipose tissue. In particular, individuals with high muscle mass may be incorrectly placed into the overweight or obese category despite having low overall body fat (Kyle et al., 2003). A more accurate anthropometric measure could be used in future studies such as body fat percentage or measures of central obesity such as visceral adiposity derived from body composition scans.

Despite limitations, BMI in children and adolescents has shown to be an accurate predictor of future health and disease outcomes (Kelishadi et al., 2015). Longitudinal studies have consistently demonstrated that higher BMI is correlated with metabolic and inflammatory diseases (Aarestrup & Koopmans, 2016; Twig et al., 2016; Zimmermann et al., 2017). Additionally, data regarding participants physical activity, sedentary behaviour and dietary habits were not collected. In particular, consumption of ultra processed “junk” foods for just 4 days was associated with the rapid development of cognitive impairment in adults (Attuquayefio et al., 2017), and dietary behaviours vary widely across individuals. As such, individuals may consume a diet replete in foods high in refined sugar and saturated fat, yet still maintain a “healthy” BMI, sometimes known as “normal weight, metabolically obese” (Oliveros et al., 2014). This definition encompasses individuals that have metabolic dysregulation as opposed to increased weight to height ratios, and supports the need for an updated definition of obesity based on adiposity, not on body weight.

Although we did not test cognitive performance in our sample, associations between obesity and impairment across multiple cognitive domains have been noted, in particular executive functioning (Yang et al., 2018), verbal memory (Cournot et al., 2006), attention (Fergenbaum et al., 2009), inhibition and decision making (Appelhans, 2009; Davis et al., 2020; Fagundo et al., 2012).

Considering that energy metabolism is significantly altered in obesity, the potential complex implications of neurometabolic dysregulation induced by increased BMI across the lifespan is important. Although physiological and lifestyle confounds may also impact neurophysiology, they underscore the significance of incorporating such information for enhanced elucidation and contextualization of the findings. This study contributes valuable neurophysiological insight into associations between functional connectivity patterns in young people with elevated BMI.

### Conclusions

Given that inhibitory signalling in the frontal cortex can influence decision-making around food, these results have important implications for both weight management and brain health in individuals in present society. This is of particular relevance as palatable energy dense foods are abundantly available, particularly to children and adolescents, and may have enduring impacts on brain function.

## Acknowledgements

ACR received funding in the form of an Investigator Award from the National Health and Medical Research Council (Australia).

## References

Aarestrup, F. M., & Koopmans, M. G. (2016). Sharing Data for Global Infectious Disease Surveillance and Outbreak Detection. Trends in Microbiology, 24(4), 241–245. 10.1016/j.tim.2016.01.009

Appelhans, B. M. (2009). Neurobehavioral Inhibition of Reward-driven Feeding: Implications for Dieting and Obesity. Obesity, 17(4), 640–647. 10.1038/oby.2008.638

Attuquayefio, T., Stevenson, R. J., Oaten, M. J., & Francis, H. M. (2017). A four-day Western-style dietary intervention causes reductions in hippocampal-dependent learning and memory and interoceptive sensitivity. PloS One, 12(2), e0172645. 10.1371/journal.pone.0172645

Babiloni, C., Marzano, N., Lizio, R., Valenzano, A., Triggiani, A. I., Petito, A., Bellomo, A., Lecce, B., Mundi, C., Soricelli, A., Limatola, C., Cibelli, G., & Del Percio, C. (2011). Resting state cortical electroencephalographic rhythms in subjects with normal and abnormal body weight. NeuroImage, 58(2), 698–707. 10.1016/j.neuroimage.2011.05.080

Baek, K., Morris, L. S., Kundu, P., & Voon, V. (2017). Disrupted resting-state brain network properties in obesity: Decreased global and putaminal cortico-striatal network efficiency. Psychological Medicine, 47(4), 585–596. 10.1017/S0033291716002646

Baker, K. D., & Reichelt, A. C. (2016). Impaired fear extinction retention and increased anxiety-like behaviours induced by limited daily access to a high-fat/high-sugar diet in male rats: Implications for diet-induced prefrontal cortex dysregulation. Neurobiology of Learning and Memory, 136, 127–138. 10.1016/j.nlm.2016.10.002

Barbosa, D. A. N., Gattas, S., Salgado, J. S., Kuijper, F. M., Wang, A. R., Huang, Y., Kakusa, B., Leuze, C., Luczak, A., Rapp, P., Malenka, R. C., Hermes, D., Miller, K. J., Heifets, B. D., Bohon, C., McNab, J. A., & Halpern, C. H. (2023). An orexigenic subnetwork within the human hippocampus. Nature, 621(7978), 381–388. 10.1038/s41586-023-06459-w

Biswal, B. B., Kylen, J. V., & Hyde, J. S. (1997). Simultaneous assessment of flow and BOLD signals in resting-state functional connectivity maps. NMR in Biomedicine, 10(4–5), 165–170. 10.1002/(SICI)1099-1492(199706/08)10:4/5<165::AID-NBM454>3.0.CO;2-7

Bocarsly, M. E., Fasolino, M., Kane, G. A., LaMarca, E. A., Kirschen, G. W., Karatsoreos, I. N., McEwen, B. S., & Gould, E. (2015). Obesity diminishes synaptic markers, alters microglial morphology, and impairs cognitive function. Proceedings of the National Academy of Sciences of the United States of America, 112(51), 15731–15736. 10.1073/pnas.1511593112

Brookes, M. J., Woolrich, M. W., & Barnes, G. R. (2012). Measuring functional connectivity in MEG: A multivariate approach insensitive to linear source leakage. NeuroImage, 63(2), 910–920. 10.1016/j.neuroimage.2012.03.048

Buckner, R. L., Andrews-Hanna, J. R., & Schacter, D. L. (2008). The brain’s default network: Anatomy, function, and relevance to disease. Annals of the New York Academy of Sciences, 1124, 1–38. 10.1196/annals.1440.011

Buzsáki, G., & Wang, X.-J. (2012). Mechanisms of Gamma Oscillations. Annual Review of Neuroscience, 35(1), 203–225. 10.1146/annurev-neuro-062111-150444

Carlén, M., Meletis, K., Siegle, J. H., Cardin, J. A., Futai, K., Vierling-Claassen, D., Rühlmann, C., Jones, S. R., Deisseroth, K., Sheng, M., Moore, C. I., & Tsai, L.-H. (2012). A critical role for NMDA receptors in parvalbumin interneurons for gamma rhythm induction and behavior. Molecular Psychiatry, 17(5), 537–548. 10.1038/mp.2011.31

Casey, B., Glatt, C. E., & Lee, F. S. (2015). Treating the Developing versus Developed Brain: Translating Preclinical Mouse and Human Studies. Neuron, 86(6), 1358–1368. 10.1016/j.neuron.2015.05.020

Cherbuin, N., Sargent-Cox, K., Fraser, M., Sachdev, P., & Anstey, K. J. (2015). Being overweight is associated with hippocampal atrophy: The PATH Through Life Study. International Journal of Obesity (2005), *39*(10), 1509–1514. 10.1038/ijo.2015.106

Clasen, P. C., Beevers, C. G., Mumford, J. A., & Schnyer, D. M. (2014). Cognitive control network connectivity in adolescent women with and without a parental history of depression. Developmental Cognitive Neuroscience, 7, 13–22. 10.1016/j.dcn.2013.10.008

Colclough, G. L., Brookes, M. J., Smith, S. M., & Woolrich, M. W. (2015). A symmetric multivariate leakage correction for MEG connectomes. NeuroImage, 117, 439–448. 10.1016/j.neuroimage.2015.03.071

Cournot, M., Marquié, J. C., Ansiau, D., Martinaud, C., Fonds, H., Ferrières, J., & Ruidavets, J. B. (2006). Relation between body mass index and cognitive function in healthy middle-aged men and women. Neurology, 67(7), 1208–1214. 10.1212/01.wnl.0000238082.13860.50

Davis, J. A., Paul, J. R., McMeekin, L. J., Nason, S. R., Antipenko, J. P., Yates, S. D., Cowell, R. M., Habegger, K. M., & Gamble, K. L. (2020). High-Fat and High-Sucrose Diets Impair Time-of-Day Differences in Spatial Working Memory of Male Mice. Obesity, 28(12), 2347–2356. 10.1002/oby.22983

de Pasquale, F., Della Penna, S., Snyder, A. Z., Marzetti, L., Pizzella, V., Romani, G. L., & Corbetta, M. (2012). A Cortical Core for Dynamic Integration of Functional Networks in the Resting Human Brain. Neuron, 74(4), 753–764. 10.1016/j.neuron.2012.03.031

Dekkers, I. A., Jansen, P. R., & Lamb, H. J. (2019). Obesity, Brain Volume, and White Matter Microstructure at MRI: A Cross-sectional UK Biobank Study. Radiology, 291(3), 763–771. 10.1148/radiol.2019181012

Del Percio, C., Triggiani, A. I., Marzano, N., Valenzano, A., De Rosas, M., Petito, A., Bellomo, A., Lecce, B., Mundi, C., Infarinato, F., Soricelli, A., Limatola, C., Cibelli, G., & Babiloni, C. (2013). Poor desynchronisation of resting-state eyes-open cortical alpha rhythms in obese subjects without eating disorders. Clinical Neurophysiology: Official Journal of the International Federation of Clinical Neurophysiology, 124(6), 1095–1105. 10.1016/j.clinph.2013.01.001

Del Pino, I., García-Frigola, C., Dehorter, N., Brotons-Mas, J. R., Alvarez-Salvado, E., Martínez de Lagrán, M., Ciceri, G., Gabaldón, M. V., Moratal, D., Dierssen, M., Canals, S., Marín, O., & Rico, B. (2013). Erbb4 deletion from fast-spiking interneurons causes schizophrenia-like phenotypes. Neuron, 79(6), 1152–1168. 10.1016/j.neuron.2013.07.010

Dingess, P. M., Harkness, J. H., Slaker, M., Zhang, Z., Wulff, S. S., Sorg, B. A., & Brown, T. E. (2018). Consumption of a High-Fat Diet Alters Perineuronal Nets in the Prefrontal Cortex. Neural Plasticity, 2018, 2108373. 10.1155/2018/2108373

Doucet, É., McInis, K., & Mahmoodianfard, S. (2018). Compensation in response to energy deficits induced by exercise or diet. Obesity Reviews: An Official Journal of the International Association for the Study of Obesity, 19 *Suppl 1*, 36–46. 10.1111/obr.12783

Evans, A. C. & Brain Development Cooperative Group. (2006). The NIH MRI study of normal brain development. NeuroImage, 30(1), 184–202. 10.1016/j.neuroimage.2005.09.068

Fagundo, A. B., de la Torre, R., Jiménez-Murcia, S., Agüera, Z., Granero, R., Tárrega, S., Botella, C., Baños, R., Fernández-Real, J. M., Rodríguez, R., Forcano, L., Frühbeck, G., Gómez-Ambrosi, J., Tinahones, F. J., Fernández-García, J. C., Casanueva, F. F., & Fernández-Aranda, F. (2012). Executive functions profile in extreme eating/weight conditions: From anorexia nervosa to obesity. PloS One, 7(8), e43382. 10.1371/journal.pone.0043382

Fransson, P., & Marrelec, G. (2008). The precuneus/posterior cingulate cortex plays a pivotal role in the default mode network: Evidence from a partial correlation network analysis. NeuroImage, 42(3), 1178–1184. 10.1016/j.neuroimage.2008.05.059

Fries, P. (2015). Rhythms for Cognition: Communication through Coherence. Neuron, 88(1), 220–235. 10.1016/j.neuron.2015.09.034

Geha, P., Cecchi, G., Todd Constable, R., Abdallah, C., & Small, D. M. (2017). Reorganization of brain connectivity in obesity. Human Brain Mapping, 38(3), 1403–1420. 10.1002/hbm.23462

Giedd, J. N. (2004). Structural magnetic resonance imaging of the adolescent brain. Ann N Y Acad Sci, 1021, 77–85. 10.1196/annals.1308.009

Gil Ávila, C., Bott, F. S., Tiemann, L., Hohn, V. D., May, E. S., Nickel, M. M., Zebhauser, P. T., Gross, J., & Ploner, M. (2023). DISCOVER-EEG: An open, fully automated EEG pipeline for biomarker discovery in clinical neuroscience. Scientific Data, 10(1), 613. 10.1038/s41597-023-02525-0

Greenwood, C. E., & Winocur, G. (2005). High-fat diets, insulin resistance and declining cognitive function. Neurobiology of Aging, 26 *Suppl 1*, 42–45. 10.1016/j.neurobiolaging.2005.08.017

Guan, A., Wang, S., Huang, A., Qiu, C., Li, Y., Li, X., Wang, J., Wang, Q., & Deng, B. (2022). The role of gamma oscillations in central nervous system diseases: Mechanism and treatment. Frontiers in Cellular Neuroscience, 16, 962957. 10.3389/fncel.2022.962957

Guillemot-Legris, O., & Muccioli, G. G. (2017). Obesity-Induced Neuroinflammation: Beyond the Hypothalamus. Trends in Neurosciences, 40(4), 237–253. 10.1016/j.tins.2017.02.005

Haltia, L. T., Viljanen, A., Parkkola, R., Kemppainen, N., Rinne, J. O., Nuutila, P., & Kaasinen, V. (2007). Brain white matter expansion in human obesity and the recovering effect of dieting. The Journal of Clinical Endocrinology and Metabolism, 92(8), 3278–3284. 10.1210/jc.2006-2495

Hanssen, K. Ø., Grødem, S., Fyhn, M., Hafting, T., Einevoll, G. T., Ness, T. V., & Halnes, G. (2023). Responses in fast-spiking interneuron firing rates to parameter variations associated with degradation of perineuronal nets. Journal of Computational Neuroscience, 51(2), 283–298. 10.1007/s10827-023-00849-9

Huang, M.-X., Huang, C. W., Harrington, D. L., Nichols, S., Robb-Swan, A., Angeles-Quinto, A., Le, L., Rimmele, C., Drake, A., Song, T., Huang, J. W., Clifford, R., Ji, Z., Cheng, C.-K., Lerman, I., Yurgil, K. A., Lee, R. R., & Baker, D. G. (2019). Marked Increases in Resting-State MEG Gamma-Band Activity in Combat-Related Mild Traumatic Brain Injury. Cerebral Cortex. 10.1093/cercor/bhz087

Hunt, B. A. E., Wong, S. M., Vandewouw, M. M., Brookes, M. J., Dunkley, B. T., Taylor, M. J., Wong, S. M., Taylor, M. J., Dunkley, B. T., Brookes, M. J., & Hunt, B. A. E. (2019). Spatial and spectral trajectories in typical neurodevelopment from childhood to middle age. Network Neuroscience, 1–63. 10.1162/netn_a_00077

Jalbrzikowski, M., Larsen, B., Hallquist, M. N., Foran, W., Calabro, F., & Luna, B. (2017). Development of White Matter Microstructure and Intrinsic Functional Connectivity Between the Amygdala and Ventromedial Prefrontal Cortex: Associations With Anxiety and Depression. Biological Psychiatry, 82(7), 511–521. 10.1016/j.biopsych.2017.01.008

Jensen, O., & Mazaheri, A. (2010). Shaping functional architecture by oscillatory alpha activity: Gating by inhibition. Frontiers in Human Neuroscience, 4, 186. 10.3389/fnhum.2010.00186

Keating, C., Backholer, K., & Peeters, A. (2014). Prevalence of overweight and obesity in children and adults. The Lancet, 384(9960), 2107–2108. 10.1016/S0140-6736(14)62367-9

Kelishadi, R., Mirmoghtadaee, P., Najafi, H., & Keikha, M. (2015). Systematic review on the association of abdominal obesity in children and adolescents with cardio-metabolic risk factors. Journal of Research in Medical Sciences: The Official Journal of Isfahan University of Medical Sciences, 20(3), 294–307.

Korotkova, T., Fuchs, E. C., Ponomarenko, A., von Engelhardt, J., & Monyer, H. (2010). NMDA receptor ablation on parvalbumin-positive interneurons impairs hippocampal synchrony, spatial representations, and working memory. Neuron, 68(3), 557–569. 10.1016/j.neuron.2010.09.017

Kösling, C., Schäfer, L., Hübner, C., Sebert, C., Hilbert, A., & Schmidt, R. (2022). Food-Induced Brain Activity in Children with Overweight or Obesity versus Normal Weight: An Electroencephalographic Pilot Study. Brain Sciences, 12(12), 1653. 10.3390/brainsci12121653

Kuczmarski, R. J., Ogden, C. L., Guo, S. S., Grummer-Strawn, L. M., Flegal, K. M., Mei, Z., Wei, R., Curtin, L. R., Roche, A. F., & Johnson, C. L. (2002). 2000 CDC Growth Charts for the United States: Methods and development. Vital and Health Statistics. Series 11, Data from the National Health Survey, 246, 1–190.

Kyle, U. G., Genton, L., Hans, D., & Pichard, C. (2003). Validation of a bioelectrical impedance analysis equation to predict appendicular skeletal muscle mass (ASMM). *Clinical Nutrition (Edinburgh*, Scotland), 22(6), 537–543. 10.1016/s0261-5614(03)00048-7

Labouesse, M. A., Lassalle, O., Richetto, J., Iafrati, J., Weber-Stadlbauer, U., Notter, T., Gschwind, T., Pujadas, L., Soriano, E., Reichelt, A. C., Labouesse, C., Langhans, W., Chavis, P., & Meyer, U. (2017). Hypervulnerability of the adolescent prefrontal cortex to nutritional stress via reelin deficiency. Molecular Psychiatry, 22(7), 961–971. 10.1038/mp.2016.193

Lensjø, K. K., Lepperød, M. E., Dick, G., Hafting, T., & Fyhn, M. (2017). Removal of Perineuronal Nets Unlocks Juvenile Plasticity Through Network Mechanisms of Decreased Inhibition and Increased Gamma Activity. The Journal of Neuroscience: The Official Journal of the Society for Neuroscience, 37(5), 1269–1283. 10.1523/JNEUROSCI.2504-16.2016

Lowe, C. J., Morton, J. B., & Reichelt, A. C. (2020). Adolescent obesity and dietary decision making-a brain-health perspective. The Lancet. Child & Adolescent Health, 4(5), 388–396. 10.1016/S2352-4642(19)30404-3

Lowe, C. J., Reichelt, A. C., & Hall, P. A. (2019). The Prefrontal Cortex and Obesity: A Health Neuroscience Perspective. Trends in Cognitive Sciences, 23(4), 349–361. 10.1016/j.tics.2019.01.005

Mahmoud, R., Kimonis, V., & Butler, M. G. (2022). Genetics of Obesity in Humans: A Clinical Review. International Journal of Molecular Sciences, 23(19), 11005. 10.3390/ijms231911005

Massar, S. A. A., Kenemans, J. L., & Schutter, D. J. L. G. (2014). Resting-state EEG theta activity and risk learning: Sensitivity to reward or punishment? International Journal of Psychophysiology, 91(3), 172–177. 10.1016/j.ijpsycho.2013.10.013

Mauney, S. A., Athanas, K. M., Pantazopoulos, H., Shaskan, N., Passeri, E., Berretta, S., & Woo, T.-U. W. (2013). Developmental pattern of perineuronal nets in the human prefrontal cortex and their deficit in schizophrenia. Biological Psychiatry, 74(6), 427–435. 10.1016/j.biopsych.2013.05.007

Menon, V. (2011). Large-scale brain networks and psychopathology: A unifying triple network model. In Trends in Cognitive Sciences (Vol. 15, Issue 10, pp. 483–506). 10.1016/j.tics.2011.08.003

Mitchell, J. A., Mattocks, C., Ness, A. R., Leary, S. D., Pate, R. R., Dowda, M., Blair, S. N., & Riddoch, C. (2009). Sedentary Behaviour and Obesity in a Large Cohort of Children. *Obesity (Silver Spring*, Md.), 17(8), 1596–1602. 10.1038/oby.2009.42

Ng, M., Fleming, T., Robinson, M., Thomson, B., Graetz, N., Margono, C., Mullany, E. C., Biryukov, S., Abbafati, C., Abera, S. F., Abraham, J. P., Abu-Rmeileh, N. M. E., Achoki, T., AlBuhairan, F. S., Alemu, Z. A., Alfonso, R., Ali, M. K., Ali, R., Guzman, N. A., … Gakidou, E. (2014). Global, regional, and national prevalence of overweight and obesity in children and adults during 1980-2013: A systematic analysis for the Global Burden of Disease Study 2013. *Lancet (London*, England), 384(9945), 766–781. 10.1016/S0140-6736(14)60460-8

Olde Dubbelink, K. T. E., Felius, A., Verbunt, J. P. A., van Dijk, B. W., Berendse, H. W., Stam, C. J., & Delemarre-van de Waal, H. A. (2008). Increased Resting-State Functional Connectivity in Obese Adolescents; A Magnetoencephalographic Pilot Study. PLoS ONE, 3(7), e2827. 10.1371/journal.pone.0002827

Oliveros, E., Somers, V. K., Sochor, O., Goel, K., & Lopez-Jimenez, F. (2014). The concept of normal weight obesity. Progress in Cardiovascular Diseases, 56(4), 426–433. 10.1016/j.pcad.2013.10.003

Oostenveld, R., Fries, P., Maris, E., & Schoffelen, J. M. (2011). FieldTrip: Open source software for advanced analysis of MEG, EEG, and invasive electrophysiological data. Computational Intelligence and Neuroscience, 2011. 10.1155/2011/156869

Orekhova, E. V., Stroganova, T. A., Schneiderman, J. F., Lundström, S., Riaz, B., Sarovic, D., Sysoeva, O. V., Brant, G., Gillberg, C., & Hadjikhani, N. (2018). Neural gain control measured through cortical gamma oscillations is associated with sensory sensitivity. Human Brain Mapping, 40(5), 1583–1593. 10.1002/hbm.24469

Pintana, H., Lietzau, G., Augestad, I. L., Chiazza, F., Nyström, T., Patrone, C., & Darsalia, V. (2019). Obesity-induced type 2 diabetes impairs neurological recovery after stroke in correlation with decreased neurogenesis and persistent atrophy of parvalbumin-positive interneurons. Clinical Science, 133(13), 1367–1386. 10.1042/CS20190180

Reichelt, A. C. (2016). Adolescent Maturational Transitions in the Prefrontal Cortex and Dopamine Signaling as a Risk Factor for the Development of Obesity and High Fat/High Sugar Diet Induced Cognitive Deficits. Frontiers in Behavioral Neuroscience, 10, 189. 10.3389/fnbeh.2016.00189

Reichelt, A. C., Gibson, G. D., Abbott, K. N., & Hare, D. J. (2019). A high-fat high-sugar diet in adolescent rats impairs social memory and alters chemical markers characteristic of atypical neuroplasticity and parvalbumin interneuron depletion in the medial prefrontal cortex. Food & Function, 10(4), 1985–1998. 10.1039/c8fo02118j

Reichelt, A. C., & Rank, M. M. (2017). The impact of junk foods on the adolescent brain. Birth Defects Research, 109(20), 1649–1658. 10.1002/bdr2.1173

Roux, F., Wibral, M., Singer, W., Aru, J., & Uhlhaas, P. J. (2013). The Phase of Thalamic Alpha Activity Modulates Cortical Gamma-Band Activity: Evidence from Resting-State MEG Recordings. Journal of Neuroscience, 33(45), 17827–17835. 10.1523/JNEUROSCI.5778-12.2013

Rubinov, M., & Sporns, O. (2010). Complex network measures of brain connectivity: Uses and interpretations. NeuroImage, 52(3), 1059–1069. 10.1016/j.neuroimage.2009.10.003

Sandoval-Salazar, C., Ramírez-Emiliano, J., Trejo-Bahena, A., Oviedo-Solís, C. I., & Solís-Ortiz, M. S. (2016). A high-fat diet decreases GABA concentration in the frontal cortex and hippocampus of rats. Biological Research, 49(1), 15. 10.1186/s40659-016-0075-6

Schmidt, R., Sebert, C., Kösling, C., Grunwald, M., Hilbert, A., Hübner, C., & Schäfer, L. (2018). Neuropsychological and Neurophysiological Indicators of General and Food-Specific Impulsivity in Children with Overweight and Obesity: A Pilot Study. Nutrients, 10(12), 1983. 10.3390/nu10121983

Seabrook, L. T., Naef, L., Baimel, C., Judge, A. K., Kenney, T., Ellis, M., Tayyab, T., Armstrong, M., Qiao, M., Floresco, S. B., & Borgland, S. L. (2023). Disinhibition of the orbitofrontal cortex biases decision-making in obesity. Nature Neuroscience, 26(1), 92–106. 10.1038/s41593-022-01210-6

Seeley, W. W. (2019). The Salience Network: A Neural System for Perceiving and Responding to Homeostatic Demands. Journal of Neuroscience, 39(50), 9878–9882. 10.1523/JNEUROSCI.1138-17.2019

Sekihara, K., Nagarajan, S. S., Poeppel, D., Marantz, A., & Miyashita, Y. (2001). Reconstructing spatio-temporal activities of neural sources using an MEG vector beamformer technique. IEEE Transactions on Bio-Medical Engineering, 48(7), 760–771. 10.1109/10.930901

Sherman, L. E., Rudie, J. D., Pfeifer, J. H., Masten, C. L., McNealy, K., & Dapretto, M. (2014). Development of the Default Mode and Central Executive Networks across early adolescence: A longitudinal study. Developmental Cognitive Neuroscience, 10, 148–159. 10.1016/j.dcn.2014.08.002

Silvers, J. A., Insel, C., Powers, A., Franz, P., Helion, C., Martin, R. E., Weber, J., Mischel, W., Casey, B. J., & Ochsner, K. N. (2017). vlPFC–vmPFC–Amygdala Interactions Underlie Age-Related Differences in Cognitive Regulation of Emotion. *Cerebral Cortex (New York*, NY), 27(7), 3502–3514. 10.1093/cercor/bhw073

Sohal, V. S., Zhang, F., Yizhar, O., & Deisseroth, K. (2009). Parvalbumin neurons and gamma rhythms enhance cortical circuit performance. Nature, 459(7247), 698–702. 10.1038/nature07991

Steward, T., Juaneda-Seguí, A., Mestre-Bach, G., Martínez-Zalacaín, I., Vilarrasa, N., Jiménez-Murcia, S., Fernández-Formoso, J. A., Veciana de las Heras, M., Custal, N., Virgili, N., Lopez-Urdiales, R., García-Ruiz-de-Gordejuela, A., Menchón, J. M., Soriano-Mas, C., & Fernandez-Aranda, F. (2019). What Difference Does it Make? Risk-Taking Behavior in Obesity after a Loss is Associated with Decreased Ventromedial Prefrontal Cortex Activity. Journal of Clinical Medicine, 8(10), 1551. 10.3390/jcm8101551

Stice, E., & Burger, K. (2019). Neural Vulnerability Factors for Obesity. Clinical Psychology Review, 68, 38–53. 10.1016/j.cpr.2018.12.002

Thompson, S. V., Hannon, B. A., An, R., & Holscher, H. D. (2017). Effects of isolated soluble fiber supplementation on body weight, glycemia, and insulinemia in adults with overweight and obesity: A systematic review and meta-analysis of randomized controlled trials. The American Journal of Clinical Nutrition, 106(6), 1514–1528. 10.3945/ajcn.117.163246

Tregellas, J. R., Wylie, K. P., Rojas, D. C., Tanabe, J., Martin, J., Kronberg, E., Cordes, D., & Cornier, M.-A. (2011). Altered Default Network Activity in Obesity. Obesity, 19(12), 2316–2321. 10.1038/oby.2011.119

Twig, G., Yaniv, G., Levine, H., Leiba, A., Goldberger, N., Derazne, E., Ben-Ami Shor, D., Tzur, D., Afek, A., Shamiss, A., Haklai, Z., & Kark, J. D. (2016). Body-Mass Index in 2.3 Million Adolescents and Cardiovascular Death in Adulthood. The New England Journal of Medicine, 374(25), 2430–2440. 10.1056/NEJMoa1503840

Tzourio-Mazoyer, N., Landeau, B., Papathanassiou, D., Crivello, F., Etard, O., Delcroix, N., Mazoyer, B., & Joliot, M. (2002). Automated anatomical labeling of activations in SPM using a macroscopic anatomical parcellation of the MNI MRI single-subject brain. NeuroImage, 15(1), 273–289. 10.1006/nimg.2001.0978

Van Veen, B. D., van Drongelen, W., Yuchtman, M., & Suzuki, a. (1997). Localization of brain electrical activity via linearly constrained minimum variance spatial filtering. IEEE Transactions on Bio-Medical Engineering, 44(9), 867–880. 10.1109/10.623056

Vanderwal, T., Kelly, C., Eilbott, J., Mayes, L. C., & Castellanos, F. X. (2015). Inscapes: A movie paradigm to improve compliance in functional magnetic resonance imaging. NeuroImage, 122, 222–232. 10.1016/j.neuroimage.2015.07.069

Vandewouw, M. M., Dunkley, B. T., Lerch, J. P., Anagnostou, E., & Taylor, M. J. (2021). Characterizing Inscapes and resting-state in MEG: Effects in typical and atypical development. NeuroImage, 225, 117524. 10.1016/j.neuroimage.2020.117524

Vega-Torres, J. D., Ontiveros-Angel, P., Terrones, E., Stuffle, E. C., Solak, S., Tyner, E., Oropeza, M., dela Peña, I., Obenaus, A., Ford, B. D., & Figueroa, J. D. (2022). Short-term exposure to an obesogenic diet during adolescence elicits anxiety-related behavior and neuroinflammation: Modulatory effects of exogenous neuregulin-1. Translational Psychiatry, 12(1), 1–20. 10.1038/s41398-022-01788-2

Vincent, J. L., Patel, G. H., Fox, M. D., Snyder, A. Z., Baker, J. T., Van Essen, D. C., Zempel, J. M., Snyder, L. H., Corbetta, M., & Raichle, M. E. (2007). Intrinsic functional architecture in the anaesthetized monkey brain. Nature, 447(7140), 83–86. 10.1038/nature05758

Wang, A., Wan, X., Zhuang, P., Jia, W., Ao, Y., Liu, X., Tian, Y., Zhu, L., Huang, Y., Yao, J., Wang, B., Wu, Y., Xu, Z., Wang, J., Yao, W., Jiao, J., & Zhang, Y. (2023). High fried food consumption impacts anxiety and depression due to lipid metabolism disturbance and neuroinflammation. Proceedings of the National Academy of Sciences, 120(18), e2221097120. 10.1073/pnas.2221097120

Wiesman, A. I., Murman, D. L., Losh, R. A., Schantell, M., Christopher-Hayes, N. J., Johnson, H. J., Willett, M. P., Wolfson, S. L., Losh, K. L., Johnson, C. M., May, P. E., & Wilson, T. W. (2022). Spatially resolved neural slowing predicts impairment and amyloid burden in Alzheimer’s disease. Brain, 145(6), 2177–2189. 10.1093/brain/awab430

Wingert, J. C., & Sorg, B. A. (2021). Impact of Perineuronal Nets on Electrophysiology of Parvalbumin Interneurons, Principal Neurons, and Brain Oscillations: A Review. Frontiers in Synaptic Neuroscience, 13, 673210. 10.3389/fnsyn.2021.673210

Womelsdorf, T., Fries, P., Mitra, P. P., & Desimone, R. (2006). Gamma-band synchronization in visual cortex predicts speed of change detection. Nature, 439(7077), 733–736. 10.1038/nature04258

Xia, M., Wang, J., & He, Y. (2013). BrainNet Viewer: A Network Visualization Tool for Human Brain Connectomics. PLoS ONE, 8(7), e68910. 10.1371/journal.pone.0068910

Yang, Y., Shields, G. S., Guo, C., & Liu, Y. (2018). Executive function performance in obesity and overweight individuals: A meta-analysis and review. Neuroscience and Biobehavioral Reviews, 84, 225–244. 10.1016/j.neubiorev.2017.11.020

Zimmermann, E., Bjerregaard, L. G., Gamborg, M., Vaag, A. A., Sørensen, T. I. A., & Baker, J. L. (2017). Childhood body mass index and development of type 2 diabetes throughout adult life-A large-scale danish cohort study. *Obesity (Silver Spring*, Md.), 25(5), 965–971. 10.1002/oby.21820

